# Spatiotemporal transcriptomics reveals distinct responses of ALDH1A1-positive and ALDH1A1-negative midbrain dopaminergic neurons to alpha-synuclein overexpression

**DOI:** 10.64898/2026.07.17.739121

**Authors:** Carmen-Jeanette Stepek, Éanna B. Ryan, Javier Villegas-Salmerón, Felicia New, Kalyan Chavda, David C. Henshall, Donato A. Di Monte, Jochen H. M. Prehn, Ayse Ulusoy, Niamh M.C. Connolly

**Author notes:** **Corresponding author:** Niamh M. C. Connolly.

## Abstract

Parkinson’s disease is characterized by the progressive and preferential degeneration of dopaminergic neurons in the *substantia nigra pars compacta*, and intraneuronal alpha-synuclein (αSyn) accumulation. Dopaminergic neurons (DANs) are anatomically and molecularly heterogeneous, but the impact of αSyn pathology on distinct subpopulations is not well defined. One midbrain DAN sub-population expresses Aldehyde Dehydrogenase 1A1 (ALDH1A1), an enzyme that detoxifies aldehyde by-products of dopamine metabolism, and has been associated with differential vulnerability. Here, we applied GeoMx spatial transcriptomics to profile ALDH1A1-positive (ALDH1A1^+^) and ALDH1A1-negative (ALDH1A1^−^) DAN subpopulations in the mouse midbrain *in situ* at 3- and 8-weeks following adeno-associated virus (AAV)-mediated αSyn overexpression. Analyzing 10,532 genes, we identified robust transcriptional differences between ALDH1A1^+^ and ALDH1A1^−^ DANs under control conditions, supporting their characterization as distinct molecular subpopulations. In AAV-αSyn-injected mice, we observed increased *Snca* expression and a reduction in ALDH1A1^−^ DANs in the ipsilateral *substantia nigra*. αSyn overexpression induced subpopulation-specific and time-dependent transcriptional responses, with dysregulation in ALDH1A1^−^ DANs characterized by early down-regulation of pathways related to synaptic function, neurotransmitter handling, and bioenergetics, including glycolysis. In contrast, ALDH1A1^+^ DANs displayed later up-regulation of genes enriched for Acetyl-CoA and cholesterol metabolism pathways, reflecting subpopulation-specific adaptations to αSyn overexpression. Analysis of human single nucleus RNA-sequencing data revealed partial conservation of the metabolic dysregulation signature. Together, our findings show that murine midbrain ALDH1A1^+^ and ALDH1A1^−^ DANs represent molecularly distinct subpopulations with divergent temporal responses to αSyn overexpression, emphasizing the importance of cell-type and disease-stage context in studies of Parkinson’s disease mechanisms.

## INTRODUCTION

Parkinson’s Disease (PD) is the second most common neurodegenerative disorder and one of the fastest growing neurological conditions worldwide^1^. Treatments are currently symptomatic, and no treatment exists to prevent, slow or cure PD, in part due to an incomplete understanding of its molecular mechanisms and pronounced disease heterogeneity^2,3^. PD is pathologically characterized by progressive dopaminergic neuron (DAN) degeneration, particularly in the ventral *substantia nigra pars compacta*, and accumulation of misfolded alpha-synuclein (αSyn) protein aggregates in Lewy bodies and neurites^4,5^. αSyn is predominantly localized to the presynaptic terminals of neurons^6,7^ where it regulates synaptic functions^8^. Its aggregation is central to PD pathogenesis^5,9^, and mutations and copy number variations in the gene encoding αSyn (*SNCA*) are causative of familial PD^10,11^.

DANs demonstrate significant molecular and functional diversity, and can be categorized by location, physiological function, electrophysiological properties, projection pattern, and gene/protein expression profile^12^. DANs also display differential vulnerability in PD, even within the midbrain. Despite the proximity of the *substantia nigra* (SN) and the ventral tegmental area (VTA), DANs in the VTA are relatively resistant to degeneration compared to DANs in the SN^13–15^. A key DAN subpopulation is characterized by expression of Aldehyde Dehydrogenase 1A1 (ALDH1A1). Within the midbrain, ALDH1A1^+^ DANs are primarily localised to the ventral tier of the SN^16–18^. ALDH1A1 is a cytosolic enzyme that catalyses the oxidation of aldehydes to carboxylic acids, and plays a key role in metabolising cytotoxic by-products of dopamine synthesis^19–21^. Inhibition of ALDH1A1 leads to increased levels of the toxic metabolite 3,4-dihydroxyphenylacetaldehyde (DOPAL), promoting toxic aggregation of αSyn^22,23^. Notably, studies report a selective loss of ALDH1A1^+^ DANs in post-mortem human PD brains and it has been postulated that the loss of ALDH1A1 mRNA and protein in these DANs may render them more susceptible to degeneration^17,24^.

Animal models over-expressing αSyn can replicate key aspects of PD pathology, including synaptic dysfunction and the selective and progressive loss of DANs in the SN^25–27^. Several models have further shown that DANs in the SN are not uniformly affected, supporting the concept that molecularly or anatomically defined DAN sub-populations differ in their response to pathological stress^13,16,28,29^. However, the relationship between ALDH1A1 expression and vulnerability remains complex. Increased vulnerability of ALDH1A1^+^ DANs has been reported in mitochondrial toxin models of PD pathology^16,30,31^, whereas transgenic mice expressing mutant αSyn demonstrated preferential vulnerability of ALDH1A1^−^ DANs, and ALDH1A1^−^ DANs also exhibited a greater burden of aggregated αSyn^17^. These findings suggest that ALDH1A1-defined DAN subpopulations may respond differently depending on the nature of the disease-associated insult, the model system, and the stage of pathology.

Novel spatial transcriptomics techniques now enable high-resolution mapping of gene expression within specific brain regions and cell types, offering an opportunity to investigate cell-type-specific responses *in situ*^32–35^. In this study, we applied GeoMx spatial transcriptomics to characterize the *in situ* molecular signatures of ALDH1A1^+^ and ALDH1A1^−^ midbrain DANs and their cell-type-specific transcriptional responses across two timepoints in a mouse model of αSyn overexpression.

## METHODS & MATERIALS

### AAV-mediated model of αSyn overexpression

This study is reported in accordance with ARRIVE guidelines. Animal experiments were approved by the Landesamt für Verbraucherschutz und Ernährung NRW (LAVE, 81-02.04.2023.A082) and complied with the German Animal Welfare Act and the EU Directive 2010/63/EU. C57BL6/NRJ mice were housed in an S2 environment at the German Centre for Neurodegenerative Diseases (DZNE) Animal Facility. All mice were housed in individually ventilated cages under specific pathogen-free conditions on a 12-hr light/dark cycle, with ad libitum access to food and water. AAV-mediated αSyn overexpression (or GFP as control) was performed using recombinant adeno-associated viral particles with a serotype 2 genome and serotype 6 capsid. The vector encoded human αSyn or GFP under the control of the human Synapsin 1 promoter, with expression enhanced by a woodchuck hepatitis virus post-transcriptional regulatory element (WPRE) and a downstream polyA signal. AAV vector production, purification, concentration, and titration were performed by Sirion Biotech, Martinsried, Germany, now part of Revvity Gene Delivery. Twelve-week-old female mice received unilateral stereotactic injections into the SN of either AAV2/6-GFP (n=7) or AAV6-SNCA (n=11) at a titer of 5×10^12^ genome copies/mL. Naïve controls (n=5) were not subjected to any injection. The sample sizes and experimental procedures were selected based on previous studies^36–38^. Animals were randomly allocated to experimental groups and, where possible, animals from different cages were distributed across groups to minimize cage-related confounding. Investigators performing surgeries were aware of group allocation, however, outcome assessment and quantitative analyses were performed blinded to treatment group where possible. Briefly, mice were anesthetized with isoflurane and placed in a stereotaxic frame. A single unilateral injection of AAV vector solution was delivered into the substantia nigra using the following coordinates relative to bregma: anteroposterior, −2.3 mm; mediolateral, ±1.1 mm; dorsoventral, −4.1 mm from the dura. A total volume of 1.5 µL was injected at a rate of 0.4 µL/min using a Hamilton syringe fitted to a glass capillary. After injection, the capillary was left in place for 5 min before slow retraction to minimize reflux. At 3- and 8-weeks post-injection, mice were sacrificed using a lethal dose of sodium pentobarbital (600 mg/kg, i.p.). All solutions used for tissue processing were prepared using nuclease free water. Mice were transcardially perfused with ice-cold saline (0.9% NaCl) from the ascending aorta and then with 4% PFA (50 mL, 10 mL/min). The brains were removed and post-fixed for another 24 hours before being transferred to sucrose for cryopreservation.

### Fixed Frozen (FxF) Tissue Preparation and Slide Mounting

Brains were post-fixed in 4% PFA, cryoprotected in graded sucrose (10%, 20%, 30% in PBS) at 4°C until equilibrated, rapidly frozen on dry ice and stored at -70°C. Coronal midbrain sections (10 μm; approx. Bregma -2.5 to -3.8 mm) were cut on a cryostat and 100 serial sections from one hemisphere (the injected side for AAV treated animals) were collected per brain. 5-6 sections per slide were mounted on Superfrost® Plus slides and stored at -70°C until use.

Following tissue processing, samples were stained for total human αSyn, as described below, and injection targeting was assessed. GFP expression was assessed by native GFP fluorescence. Only samples showing complete transgene expression in the SN were included in subsequent analyses.

### Immunofluorescence staining (TH, ALDH1A1)

Slides were brought to room temperature (RT) and washed in PBS to remove OCT. Sections were baked onto slides at 60°C to improve adherence. Tissue was then dehydrated through a series of ethanol washes (50%, 70%, 100%), followed by heat-induced antigen retrieval. Blocking in 5% normal horse serum (NHS) with 0.1% Tween-20 was performed for 30 minutes at RT. Primary antibodies against ALDH1A1 (rabbit, 1:500, Sigma HPA002123), tyrosine hydroxylase (TH; chicken, 1:500, Aves Las TYH), and total α-Synuclein (Syn211; mouse, Millipore 36-008) were applied in 1% NHS/0.1% Tween-20 and incubated overnight at 4°C. Slides were then washed and incubated in species-appropriate Alexa Fluor secondary antibodies for 1 hour at RT, washed, and coverslipped in DAPI-containing mounting medium.

### Quantification of TH^+^, ALDH1A1^+^ and ALDH1A1^−^ dopaminergic neurons

IF-labelled slides were scanned using a Zeiss AxioScan Z1 and the SN and VTA were delineated in each section. ALDH1A1^+^ and ALDH1A1^−^ DANs within regions of interest (ROIs) were manually quantified in QuPath. Nuclei in each ROI were segmented and quantified using the automated StarDist nucleus detection extension. TH^+^, TH^+^/ALDH1A1^+^ and TH^+^/ALDH1A1^−^ counts were normalized to the number of nuclei per ROI. Data analysis and visualizations were performed in R (readxl, tidyr, FSA, ggplot2, ggsignif, ggpubr, naniar). Statistical comparisons among treatment groups were conducted using the Kruskal-Wallis test followed by Dunn’s post-hoc test to compare mean ranks.

### GeoMx Spatial Transcriptomics

#### Slide Preparation

Slide preparation leading up to and including heated antigen retrieval were as described above. To expose RNA targets, sections were incubated in 1 µg/mL working solution of proteinase K, heated to 37°C, then washed with PBS. In situ hybridization was performed as directed by the standard GeoMx Slide Preparation User Manual (MAN-10087-04 for software v1.5).

Sections were blocked in GeoMx Buffer W for 30 min at RT in a humidified, light protected chamber, then incubated for 1 hour with ALDH1A1 (rabbit, 1:500, Sigma HPA002123) and tyrosine hydroxylase (TH; chicken, 1:500, Aves Las TYH) diluted in Buffer W. After washes with 2xSSC, sections were incubated for 1 hour at RT with Alexa Fluor secondary antibodies (anti-rabbit 647, anti-chicken 594; 1:500) and SYTO83 nuclear stain (1:500), washed again with 2x SSC and stored in fresh 2x SSC at 4°C until loading into the GeoMx platform.

#### Cell type segmentation, probe collection, sequencing

Slides were scanned at 20× magnification on the GeoMx Digital Spatial Profiler (DSP). IF labeling was used to delineate ROIs within the medial and lateral midbrain, encompassing the SN and VTA. ROIs were exported to ImageJ, where a custom macro enabled the manual segmentation of the border of each TH^+^/ALDH1A1^+^ and TH^+^/ALDH1A1^−^ population and generated subtype-specific masks. The resulting masks [areas of illumination (AOIs)] were imported back into the GeoMx software to guide probe collection. Probes were aspirated from AOIs into 96-well plates, and plates were incubated overnight at RT. Libraries were prepared from pooled AOIs and sequenced at the UCD Genomics Core Facility at a depth of up to 400 million reads with an Illumina NextSeq500/550.

#### QC and Data Pre-processing

Raw sequencing files (BCL format) were converted to FASTQ and then to digital count conversion (DCC) files, containing probe counts and sequencing quality metadata. Gene-level counts were calculated as the mean number of probes per AOI. The limit of quantification (LOQ) was determined per gene per AOI using:

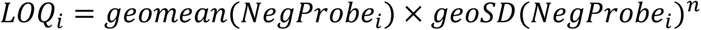

Initial pre-processing included filtering AOIs based on negative control probe counts and sequencing quality, removing 8 of 88 total AOIs. Low-performing genes, detected in <10% of AOIs or below the LOQ, were also removed, resulting in a final dataset of 10,532 genes across 80 AOIs (Table 1). Raw counts were normalized using Q3 normalization (upper quartile counts vs. negative control probe means). Principal Component Analysis, Uniform Manifold Approximation and Projection, and t-distributed Stochastic Neighbor Embedding (R packages: umap, Rtsne, standR, psych) revealed a slide-specific batch effect unrelated to biological variables. Batch correction was performed by the NanoString spatial data analysis team using a mixed-effects model with log_2_ Q3 expression as the dependent variable, incorporating batch as a random effect^39^.

**Table 1:** Number of animals, areas of interest (AOIs) and cells for each treatment group in the GeoMx® spatial transcriptomics experiment, following QC.

| Condition | # Animals | Cell type | # AOIs | # Cells | # SN AOIs | # SN cells |
| --- | --- | --- | --- | --- | --- | --- |
| naïve | 5 | ALDH1A1<sup>+</sup> | 11 | 348 | 11 | 348 |
|  |  | ALDH1A1<sup>-</sup> | 10 | 240 | 8 | 195 |
| GFP 8w | 7 | ALDH1A1<sup>+</sup> | 11 | 442 | 11 | 442 |
|  |  | ALDH1A1<sup>-</sup> | 12 | 293 | 9 | 145 |
| αSyn 3w | 5 | ALDH1A1<sup>+</sup> | 9 | 330 | 6 | 219 |
|  |  | ALDH1A1<sup>-</sup> | 8 | 210 | 5 | 95 |
| αSyn 8w | 6 | ALDH1A1<sup>+</sup> | 10 | 316 | 8 | 285 |
|  |  | ALDH1A1<sup>-</sup> | 9 | 187 | 4 | 64 |

### Differential expression and pathway enrichment analysis

The individual mouse was the experimental unit, and multiple AOIs collected from the same animal were treated as spatial replicates. Data analysis was performed in R (v 4.3.3, BiocManager 3.18). Differential expression was assessed using linear mixed models (LMMs) implemented in the NanoString Bioconductor workflow, with treatment group and DAN subtype as fixed effects and Section ID and Slide ID as random intercepts, accounting for the non-independence of multiple AOIs derived from the same animal. P-values were adjusted via Benjamini-Hochberg FDR correction.

To compare ALDH1A1+ and ALDH1A1– DANs, we fitted a LMM with subtype as the main effect and random slope by section and slide:

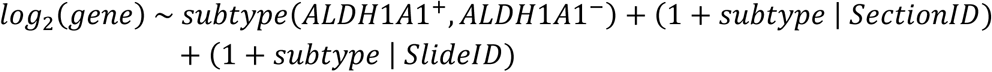

For SN-specific analyses, AOIs with <50% SN tissue were excluded. The proportion of SN tissue per AOI was calculated by anatomically delineating the SN and VTA on each GeoMx image and quantifying the number of cells within the SN region of interest relative to the total (SN + VTA) in that AOI.

To identify differential expression induced by αSyn overexpression, a temporal and treatment-specific LMM was applied to 62 SN-enriched AOIs:

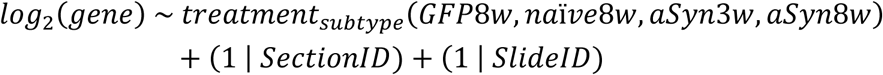

For αSyn-specific effects, we considered only genes significantly dysregulated in both αSyn8w vs. naïve8w and αSyn8w vs. GFP8w comparisons. We reasoned that DEGs in the ‘αSyn8w vs. naïve8w’ comparison only may be due to the intranigral injections and/or AAV expression, while DEGs in the ‘αSyn8w vs. GFP8w’ comparison only, may be due to GFP expression. In agreement with this, several up-regulated DEGs unique to the ‘αSyn8w vs. naïve8w’ comparison related to inflammatory pathways in both cell types (Supp. Figure 1; grey labelled genes). We note that fold changes of DEGs were strongly correlated between the two comparisons, further supporting our approach (Supp. Figure 1; purple regression line).

To identify ‘early responding genes’ (ERGs) and ‘late responding genes’ (LRGs) we categorized 8w DEGs as ERGs or LRGs by comparing their expression between the αSyn3w and αSyn8w groups:

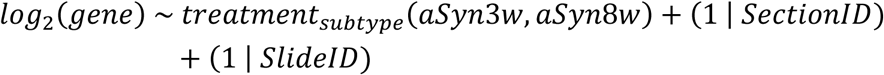

- Early responding genes (ERGs) – dysregulated at 8 weeks (αSyn8w vs controls) and statistically unchanged between 3 and 8 weeks (αSyn3w vs αSyn8w).
- Late responding genes (LRGs) – dysregulated at 8 weeks (αSyn8w vs controls) and also dysregulated between timepoints (αSyn3w vs αSyn8w). i.e., dysregulation of these genes only appeared at 8 weeks.

Pathway enrichment was performed on Gene Ontology Biological Processes (GOBP) using the enrichGO function (org.Mm.eg.db) from clusterProfiler40.

### Analysis of human snRNASeq data

Single-nuclei RNA-seq data generated by Kamath *et al.*^34^, were retrieved from GEO (GSE178265) and processed in R (v4.4.3), including control and PD SNpc post-mortem samples. Lewy Body Dementia samples were excluded. DAN clusters defined in the original study were grouped into ALDH1A1^high^ and ALDH1A1^low^ subpopulations based on ALDH1A1 expression across the 10 annotated DAN clusters. Pseudobulk count matrices were generated by aggregating nuclei by donor, disease status, and ALDH1A1 subpopulation. Differential expression analysis between PD and control samples was performed separately within ALDH1A1^high^ and ALDH1A1^low^ DANs using DESeq2^41^, with significance thresholds of |log_2_FC|>0.5 and Benjamini-Hochberg adjusted p-value <0.05.

### Data availability

FASTQ files from GeoMx were deposited on GEO under accession number GSE333766. Outputs from differential expression and pathway enrichment analyses are provided in Supp. Data 1&2.

## RESULTS

### Reduction of ALDH1A1^−^ dopaminergic neuron profiles in the substantia nigra following αSyn over-expression

To investigate differences in ALDH1A1^+^ and ALDH1A1^−^ DANs and their responses to αSyn, we utilized a well-characterized mouse model of AAV-mediated αSyn overexpression^42^. Samples were collected at 3- and 8-weeks post-injection. We first confirmed ipsilateral αSyn overexpression in the somas of TH^+^ DANs in AAV-*SNCA*-injected animals (Figure 1A). We then characterized the distribution of ALDH1A1^+^ and ALDH1A1^−^ DAN subtypes across the midbrain (Figure 1B, E). ALDH1A1^+^ DANs were predominantly distributed in the ventral tier of the SN and the VTA, while ALDH1A1^−^DANs were localized to the dorsal tier of the SN and across the VTA. This aligns with previous reports^16,31,43^.

**Figure 1:**
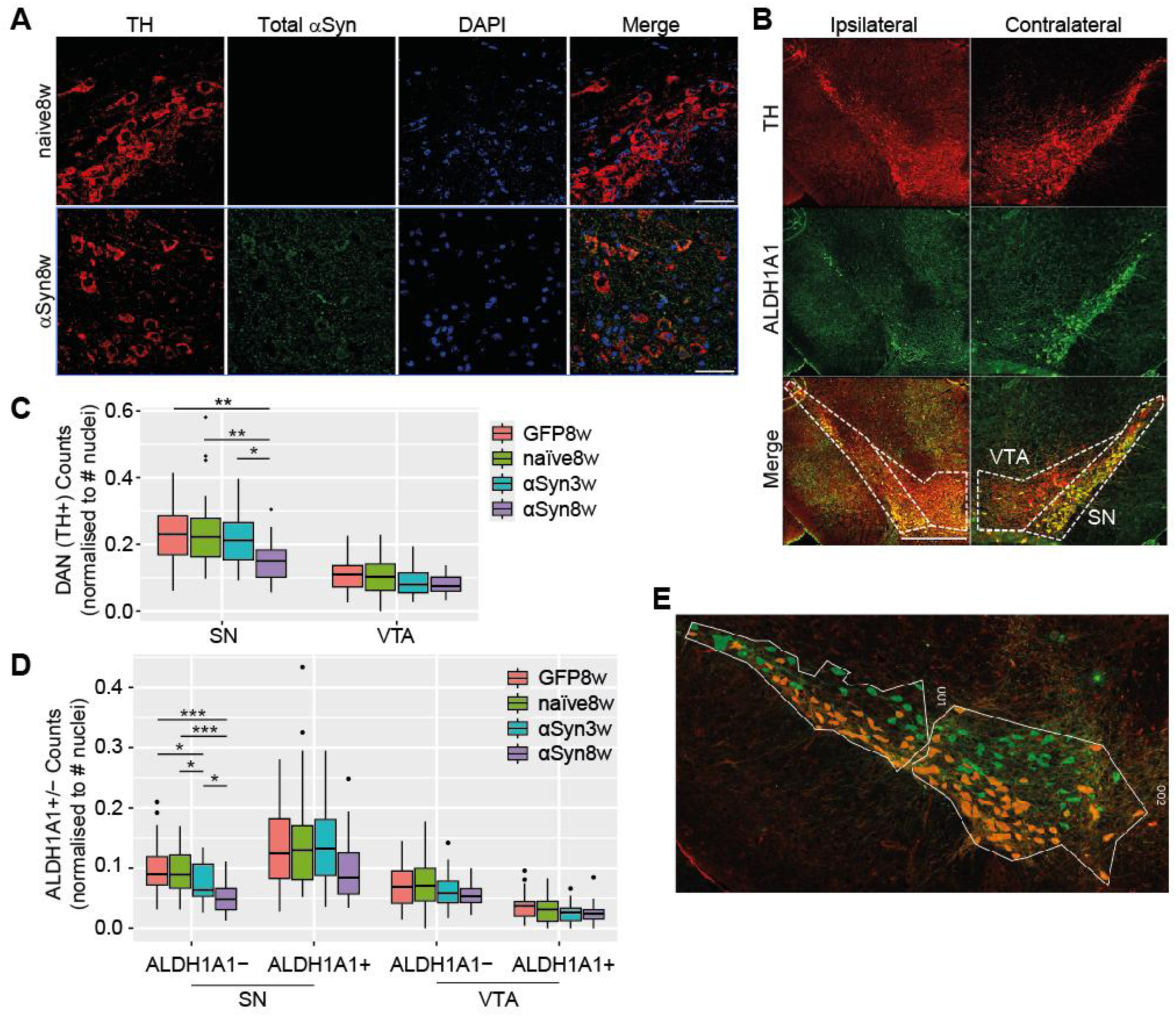
Distribution and quantification of ALDH1A1^+^ and ALDH1A1^−^ midbrain DANs in a mouse model of αSyn over-expression. (A) Immunofluorescence images of tyrosine hydroxylase (TH, red), total αSyn (green), and DAPI (blue) in coronal midbrain sections from naïve (naïve8w) and AAV-SNCA mice 8 weeks following AAV injection (αSyn8w). Scale bar 50 μm. (B) Immunofluorescence images of TH (red) and ALDH1A1 (green) in the ipsilateral and contralateral sides of an αSyn overexpressing mouse midbrain 8 weeks post-injection (αSyn8w). White dashed lines delineate the SN and VTA. Scale bar 500 μm. (C-D) Counts of C) TH^+^ neurons (normalized to # nuclei) and D) ALDH1A1^+^ and ALDH1A^−^ DANs (normalized to # nuclei) in the SN and VTA of naïve (naïve8w), AAV-GFP (GFP8w) and AAV-SNCA mice 3- and 8-weeks following AAV injection (αSyn3w, αSyn8w). Kruskal-Wallis test followed by Dunn’s post-hoc test; * p-adj< 0.05; ** p-adj<0.01; *** p-adj< 0.001. Box-plot whiskers extend to the lowest/highest value within 1.5*IQR (inter-quartile range). Data points outside these ranges are plotted as dots. (E) Representative GeoMx image from ipsilateral midbrain showing immunofluorescent TH and ALDH1A1 labeling overlaid with segmentation masks for ALDH1A1^+^ (orange) and ALDH1A1^−^ (green) DANs.

Visual inspection suggested reduced TH^+^ DANs in the ipsilateral midbrain 8 weeks following αSyn overexpression (Figure 1B). Consistent with this, we observed significant reduction of TH^+^ DANs in the ipsilateral SN at the 8-week timepoint, with no significant loss in the VTA (Figure 1C). We quantified the number of ALDH1A1^+^ and ALDH1A1^−^ DANs across the time-course of αSyn overexpression and found a significant reduction of ALDH1A1^−^ DANs in the ipsilateral SN at both 3 and 8 weeks (Figure 1D). In contrast, we did not observe a significant reduction of ALDH1A1^+^ DANs in either region, although there was a trend towards a decrease at 8 weeks in the SN (Figure 1D). No reduction in DANs was observed in GFP8w animals, indicating that the changes in αSyn-overexpressing animals were unlikely to be explained by the injection procedure itself or by AAV-induced toxicity. This suggested that ALDH1A1-defined DAN subpopulations are differentially affected in this model, with ALDH1A1^−^ DANs showing earlier and more pronounced reductions in the SN following αSyn overexpression.

### Segmentation of ALDH1A1^+^ and ALDH1A1^−^ midbrain dopaminergic neurons in situ enables cell-type-specific spatial transcriptomics

To characterize the molecular profiles of ALDH1A1^+^ and ALDH1A1^−^ midbrain DANs, we performed *in situ* spatial transcriptomics of segmented DANs from control (both naïve and GFP-injected) and αSyn over-expressing mice. We delineated ROIs spanning the medial and lateral midbrain, incorporating both the SN and VTA. TH and ALDH1A1 immunofluorescence was used to segment DAN subpopulations (Figure 1E). Barcoded hybridization probes were extracted from these segmented subpopulations to obtain spatially-resolved whole transcriptome profiles of ALDH1A1^+^ and ALDH1A1^−^ DANs *in situ*. Following quality control and batch correction to harmonise differences between slides, we retained expression data for 10,532 genes in 80 areas of interest from 23 animals (n≥5 per condition; Table 1).

Dimensionality reduction revealed separation between controls and αSyn-overexpressing samples – indicating αSyn-induced dysregulation, as well as between ALDH1A1^+^ and ALDH1A1^−^ clusters – indicating subpopulation-specific transcriptomes (Figure 2A). Notably, some αSyn samples, predominantly from the 3-week time-point, clustered closer to controls, suggesting that early-stage transcriptional changes may be more subtle.

**Figure 2:**
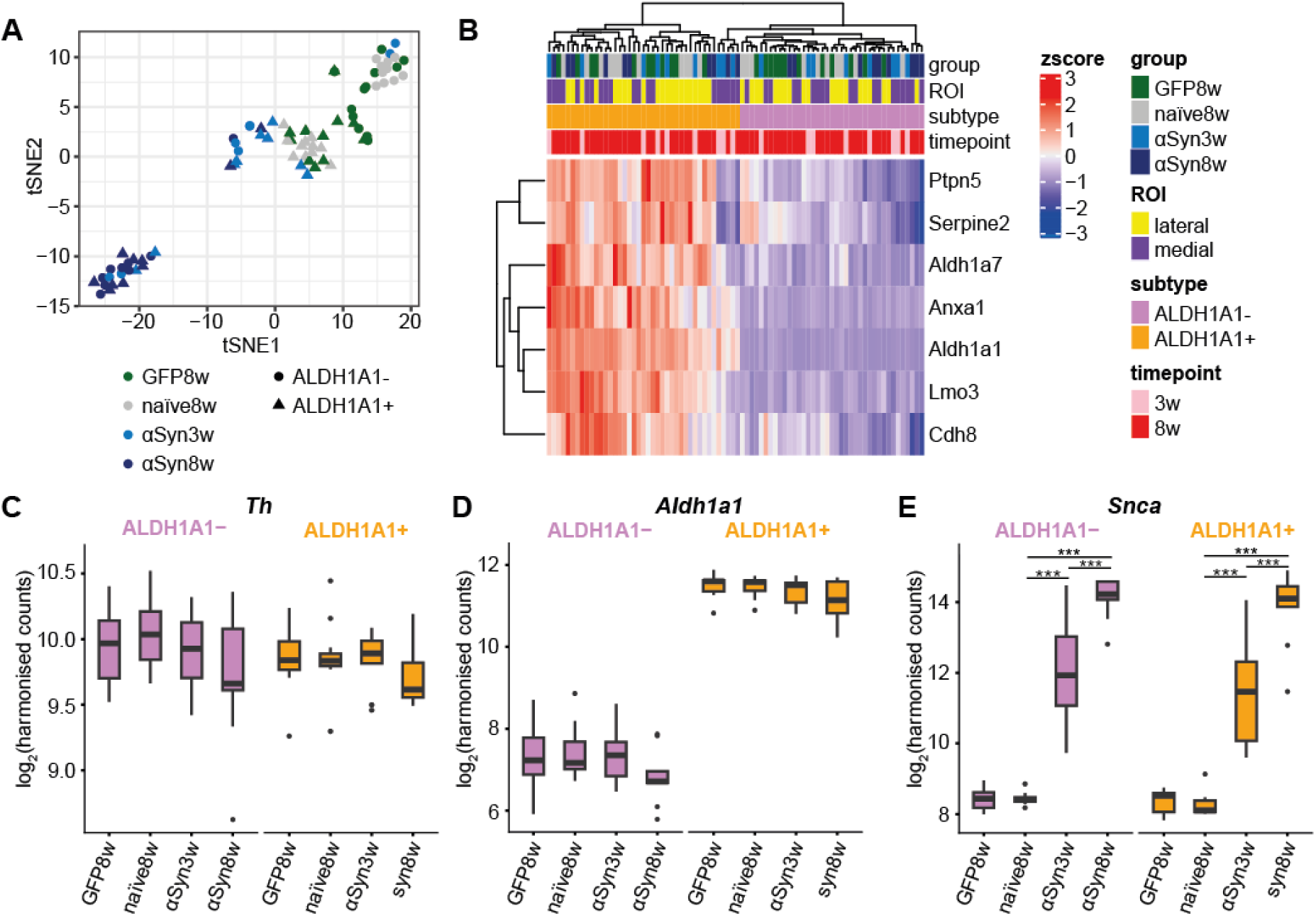
Characterisation of spatially-resolved transcriptomes collected from midbrain ALDH1A1^+^ and ALDH1A1^−^ DANs in situ. (A) Dimension reduction by t-distributed stochastic neighbour embedding (tSNE) on all samples following quality control. (B) Hierarchical clustering of previously reported ALDH1A1^+^ DAN markers. (C-E) mRNA expression levels (log-transformed counts after batch correction/harmonisation) of C) Th, D) Aldh1a1, and E) Snca, from ALDH1A1^+^ and ALDH1A1^−^ DAN samples. Linear mixed model; *** p<0.001. Box-plot whiskers extend to the lowest/highest value within 1.5*IQR (inter-quartile range). Data points outside these ranges are plotted as dots.

To verify the accurate segmentation of ALDH1A1^+^ and ALDH1A1^−^ DANs, we analysed the expression levels of specific mRNAs. Several genes have been shown to correlate with *Aldh1a1* expression in the mouse SN (*Aldh1a7*, *Anxa1, Cdh8*, *Lmo3*, *Ptpn5, Serpine2*)^44–46^, and we verified higher expression of these markers in ALDH1A1^+^ samples (Figure 2B). As expected, we also detected consistent levels of the dopaminergic marker *Th* across all samples (Figure 2C) and higher levels of *Aldh1a1* in ALDH1A1^+^ samples (Figure 2D). In addition to overexpression of total αSyn protein (Figure 1A), we found elevated levels of *Snca* mRNA in both ALDH1A1^+^ and ALDH1A1^−^ DANs in αSyn overexpressing mice, further validating the model (Figure 2E). *Snca* expression significantly increased between the 3- and 8-week timepoints. Together, this affirmed successful segmentation and extraction of the cell-type-specific transcriptome of ALDH1A1^+^ and ALDH1A1^−^ DANs in the midbrain.

### ALDH1A1^+^ and ALDH1A1^−^ DANs have distinct transcriptomic profiles, with ALDH1A1^−^ marker genes associated with synaptic functions

We next performed differential expression analysis between ALDH1A1^+^ and ALDH1A1^−^ DANs in naïve tissue, to identify transcriptomic and possibly functional differences between the subpopulations. We used a linear-mixed effects model (LMM) with stringent thresholds (|log_2_FC| ≥ 0.5, p-value <0.01), and identified 74 differentially expressed genes (DEGs; Figure 3A, Supp. Data 1a). 45 genes were more highly expressed in ALDH1A1^+^ DANs, with 29 more highly expressed in ALDH1A1^−^ DANs, defining a baseline ALDH1A1^+^/ALDH1A1^−^ transcriptomic signature (Figure 3B). Pathway enrichment analysis on these marker genes identified that ALDH1A1^+^ genes were enriched for aldehyde and fructose metabolic processes (due to expression of *Aldh1a1* and *Aldh1a7*; Figure 3C), and ALDH1A1^−^ genes were enriched for neurodevelopment, neurotransmitter activity and synaptic function (Figure 3C, Supp. Data 1b).

**Figure 3:**
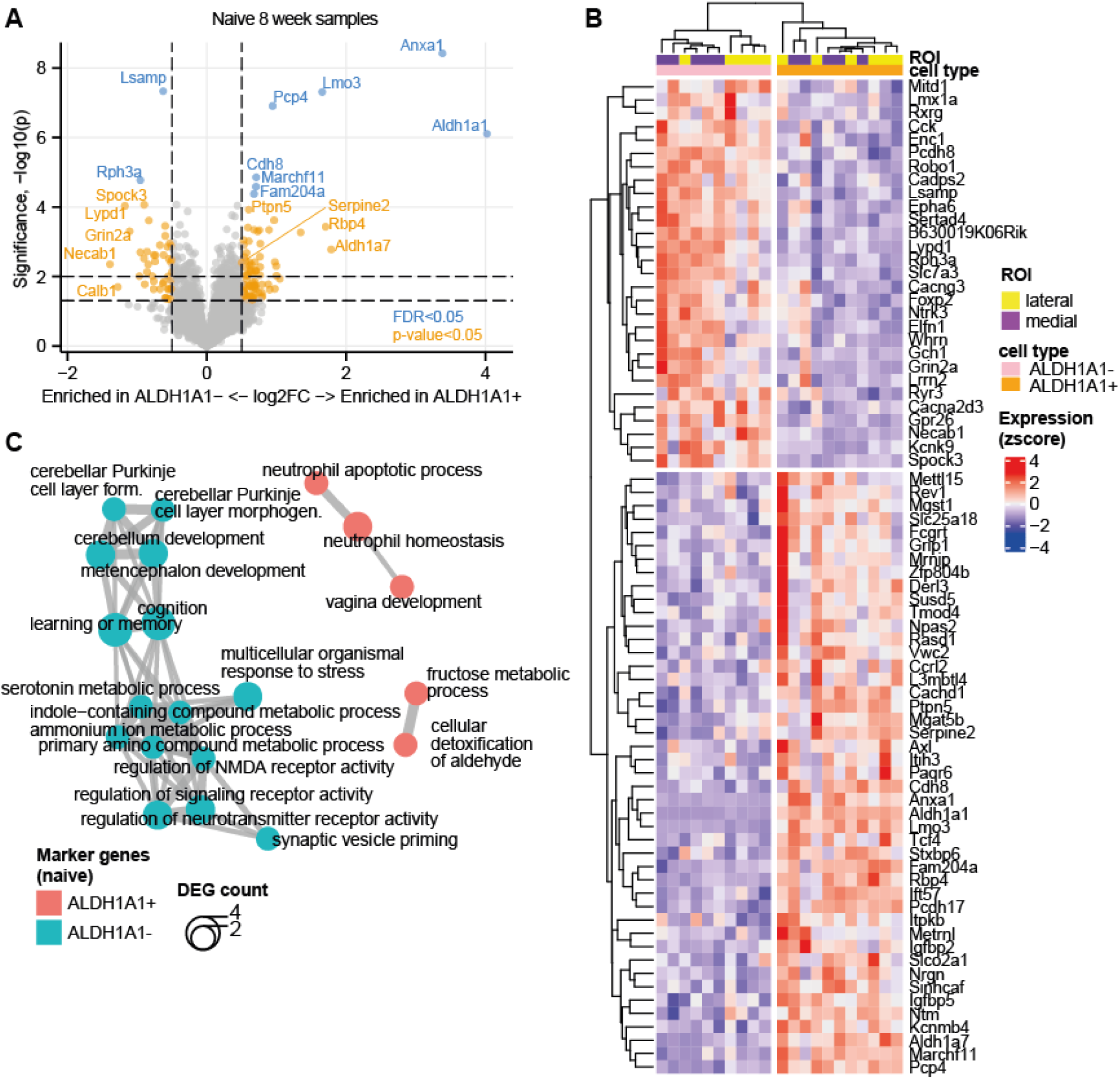
Differential expression analysis identifies a robust ALDH1A1-defined DAN signature in naïve midbrain. (A) Volcano plot showing differentially expressed genes (DEGs) between ALDH1A1^+^ and ALDH1A1^−^ DANs in naïve midbrain. Horizontal dashed lines indicate -log_10_p thresholds of 1.3 (p=0.05) and 2 (p=0.01). Vertical dashed lines indicate log_2_FC = 0.5. (B) Hierarchal clustering of z-score-normalized expression for 74 DEGs (p<0.01, |log_2_FC| > 0.5) representing an ALDH1A1-defined DAN transcriptomic signature. (C) Top 15 enriched Gene Ontology Biological Processes for marker genes from ALDH1A1^+^ and ALDH1A1^−^ DANs in naïve midbrain. Some terms have been abbreviated / removed for visualization.

### αSyn overexpression selectively impairs synaptic vesicle, neurotransmitter and bioenergetic functions in ALDH1A1^−^ DANs

We next wanted to explore how αSyn overexpression affects ALDH1A1-defined DANs, and how it may contribute to differential vulnerability. We focussed on AOIs located predominantly within the most vulnerable region of the midbrain, the SN (>50%; Figure 4A). This excluded a subset of AOIs expressing higher levels of VTA marker genes and lower levels of SN marker genes (Figure 4B). In this SN-specific analysis, we compared αSyn-overexpressing samples at 8 weeks (αSyn8w) with both naïve and GFP8w controls, and retained genes significantly dysregulated in both comparisons to ensure robustness (p<0.05 & |logFC|>0.25; see Methods).

**Figure 4.**
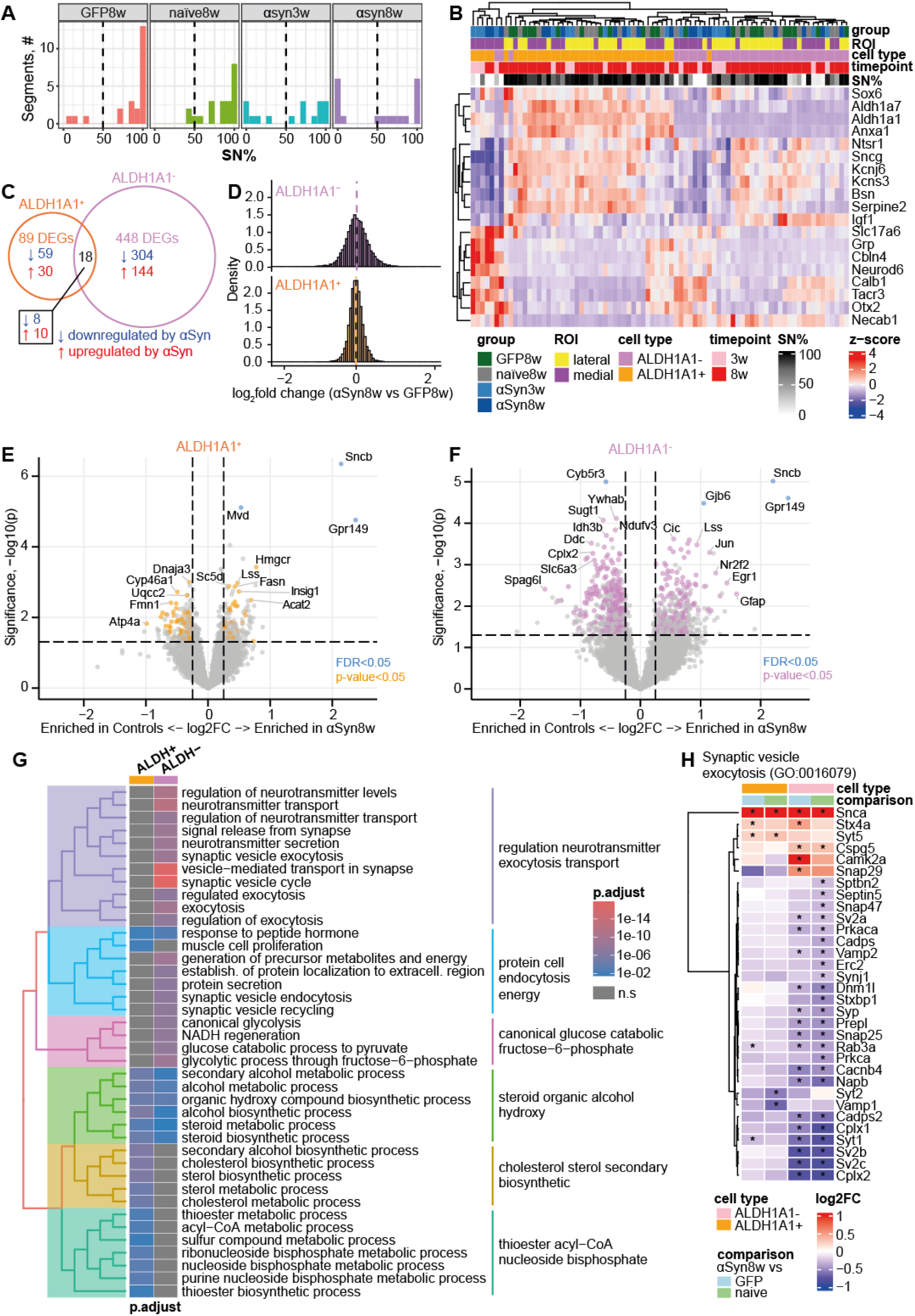
Analysis of the impact of αSyn overexpression (8w) on ALDH1A1^+^ and ALDH1A1^−^ SN dopaminergic neurons. (A) Distribution of the percentage of SN tissue (SN%) across all AOIs, stratified by treatment group, illustrating variable SN/VTA composition across sampled regions. Samples with <50% SN coverage were removed from the SN-specific analysis. (B) Hierarchical clustering of z-score normalised expression of canonical SN and VTA marker genes across all AOIs. AOIs annotated as low SN (SN% < 50%) expressed lower levels of SN markers (upper genes) and higher levels of VTA markers (lower genes). (C) Venn diagram illustrating the number of differentially expressed genes (DEGs) in ALDH1A1^+^ and ALDH1A1^−^DANs (αSyn8w vs controls). (D) Density plots of log2 fold-change (logFC) values for DEGs in each subtype (αSyn8w versus controls). (E-F) Volcano plots of DEGs for E) ALDH1A1^+^ and F) ALDH1A1^−^ DANs, for αSyn8w vs GFP8w. DEGs identified in both control comparisons (αSyn8w vs naïve and αSyn8w vs GFP8w) are coloured. Snca and EGFP (Ilog2FCI>5) have been removed for visualisation. (G) Clustered Gene Ontology Biological Process (GOBP) enrichment tree for the top 20 pathways identified from DEGs dysregulated by αSyn in ALDH1A1^+^ and ALDH1A1^−^ DANs. Terms for clusters are generated from highest frequency words within the GO term descriptions ^40^. (H) Heatmap showing log2fold change (log2FC) of genes involved in ‘Synaptic vesicle exocytosis’ (GO: 0016079). * p<0.05 in the comparisons indicated (αSyn8w vs GFP8w; αSyn8w vs naive). Only genes with p<0.05 in at least one comparison are shown.

αSyn-induced dysregulation was markedly more pronounced in SN ALDH1A1^−^ DANs (448 DEGs) compared to ALDH1A1^+^ (89 DEGs), indicating more transcriptional dysregulation in this subpopulation (Figure 4C). The magnitude of change (FC) was also greater in ALDH1A1^−^ DANs (Figure 4D). Only 18 genes were differentially expressed in both subtypes, indicating a predominantly celltype-specific response to αSyn overexpression. Shared DEGs included the highly up-regulated *Snca*, *Sncb* (α-and β-synucleins), *Gpr149* (G Protein-coupled Receptor 149), and the down-regulated nicotinic acetylcholine receptor modulator *Lynx1* (Ly6/neurotoxin 1) and synaptic vesicle regulator *Rab3c* (Ras-related protein Rab-3C). DEGs for both cell types are illustrated in Figures 4E&F, and listed in Supp. Data 2a.

Pathway enrichment of subpopulation-specific DEGs revealed divergent functional consequences of αSyn overexpression. ALDH1A1^+^ DANs were selectively enriched for sterol, specifically cholesterol, and acyl-CoA metabolism. In contrast, ALDH1A1^−^DANs showed a marked disruption of synaptic vesicle cycling, neurotransmitter processes including cathecholamine/dopamine metabolism, transport and secretion, and bioenergetic pathways including glycolysis (Figure 4G, Supp. Data 2b). Looking at synaptic vesicle exocytosis genes as an example, the selective down-regulation in ALDH1A1^−^ DANs is clear (Figure 4H), including ALDH1A1^−^-specific down-regulation of all synaptic vesicle glycoprotein 2 (Sv2) isoforms (*Sv2a*, *Sv2b*, *Sv2c*). Notably, *Sv2c* has been identified as a genetic risk factor for PD^47^.

### αSyn overexpression selectively increases sterol and cholesterol metabolism in ALDH1A1^+^ DANs at later timepoints

To delineate the temporal progression of αSyn-induced dysregulation, we categorized the αSyn8w DEGs as “early-” or “late-” responding genes by comparing their expression between the αSyn3w and αSyn8w groups (p<0.05, see Methods; Figure 5A). This revealed distinct early and late expression patterns and cellular mechanisms in ALDH1A1^+^ and ALDH1A1^−^ DANs (Figure 5B-C; Supp. Data 2a). Notably, the disruption of synaptic vesicle cycling and glycolytic pathways in ALDH1A1^−^ DANs was already evident at the 3-week timepoint, while sterol and acetyl/acyl-CoA metabolism in ALDH1A1^+^ DANs only emerged at the later 8-week timepoint (Figure 5D; Supp. Data 2c).

**Figure 5.**
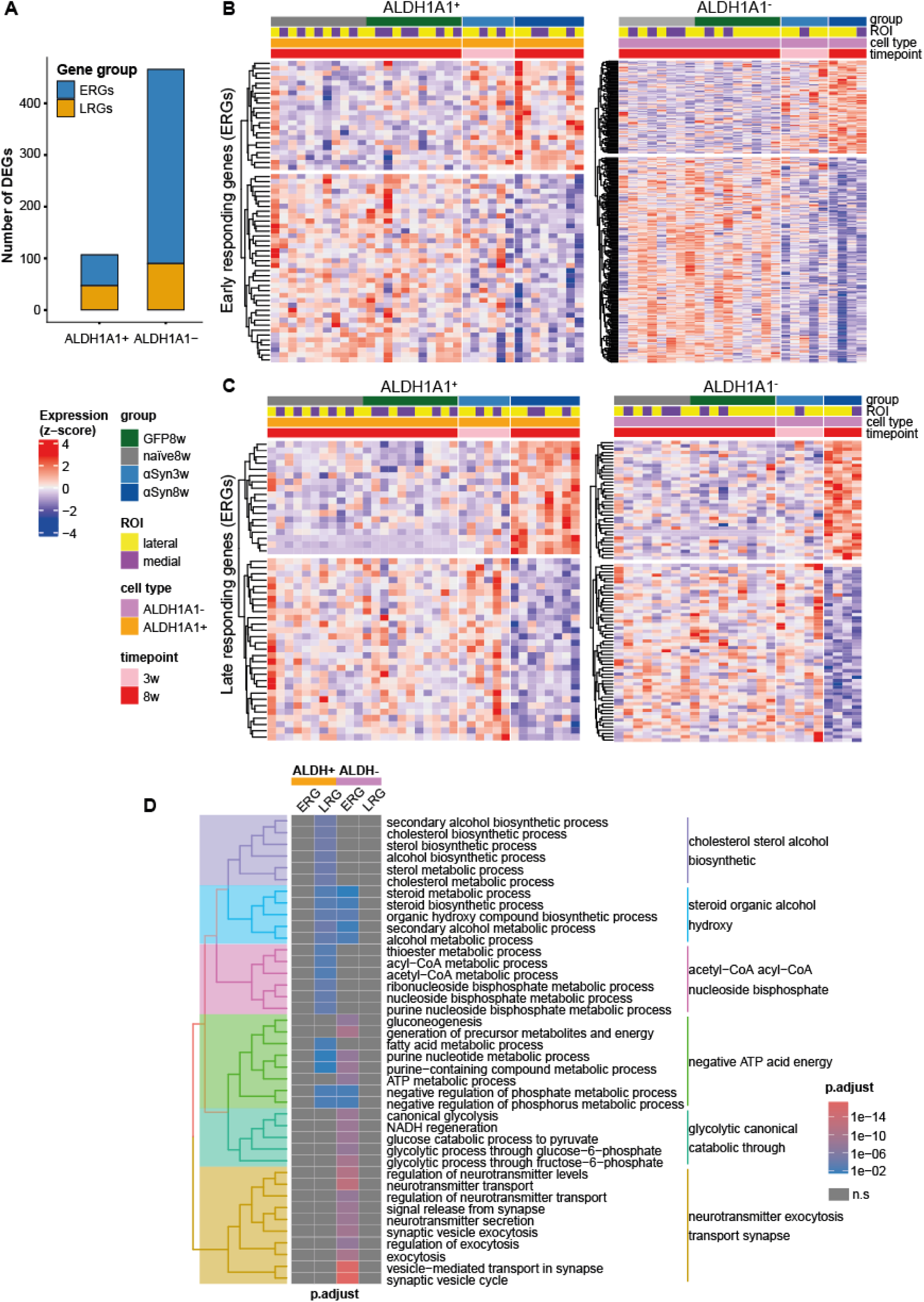
Early versus late responses to αSyn overexpression in ALDH1A1^+^ and ALDH1A1^−^ DAN subpopulations. (A) Stacked bar plots showing the number of ‘early responding genes’ (ERGs, blue) and ‘late responding genes’ (LRGs, yellow). ALDH1A1^−^ DANs: 376 ERGs, 90 LRGs. ALDH1A1^+^ DANs: 60 ERGs, 47 LRGs. (B-C) Heatmaps of z-score normalised expression for B) ERGs and C) LRGs in ALDH1A1^+^ (left) and ALDH1A1^−^ (right) SN DANs across all treatment groups (naïve8w, GFP8w, αSyn3w, αSyn8w). (D) Clustered Gene Ontology Biological Process (GOBP) enrichment tree for the top 20 pathways identified from ERGs and LRGs in ALDH1A1^+^ and ALDH1A1^−^ DANs. Terms for clusters are generated from highest frequency words within the GO term descriptions^40^. No significantly enriched pathways were identified for ALDH1A1^+^ ERGs or ALDH1A1^−^ LRGs.

Looking at individual gene expression, we found that glycolysis genes were significantly down-regulated in ALDH1A1^−^ DANs across the entire pathway (Figure 6A, B). We note that ALDH1A1^−^ ERGs were also enriched for the tricarboxylic acid (TCA) cycle and oxidative phosphorylation (Supp. Data 2c), including down-regulation of various Complex I sub-units (*Nduf5b*, *Ndufb8*, *Ndufc1*, *Ndufs2*, *Ndufv1*, *Ndufv1*), suggesting a global suppression of ATP-generating pathways in ALDH1A1^−^ DANs, even at relatively early timepoints. Interestingly, the transcription factor *Foxk1*, which promotes aerobic glycolysis and represses mitochondrial respiration^48^, was paradoxically up-regulated in ALDH1A1^−^ DANs (Figure 6A, B), potentially pointing towards an insufficient attempt to maintain glycolytic flux.

**Figure 6.**
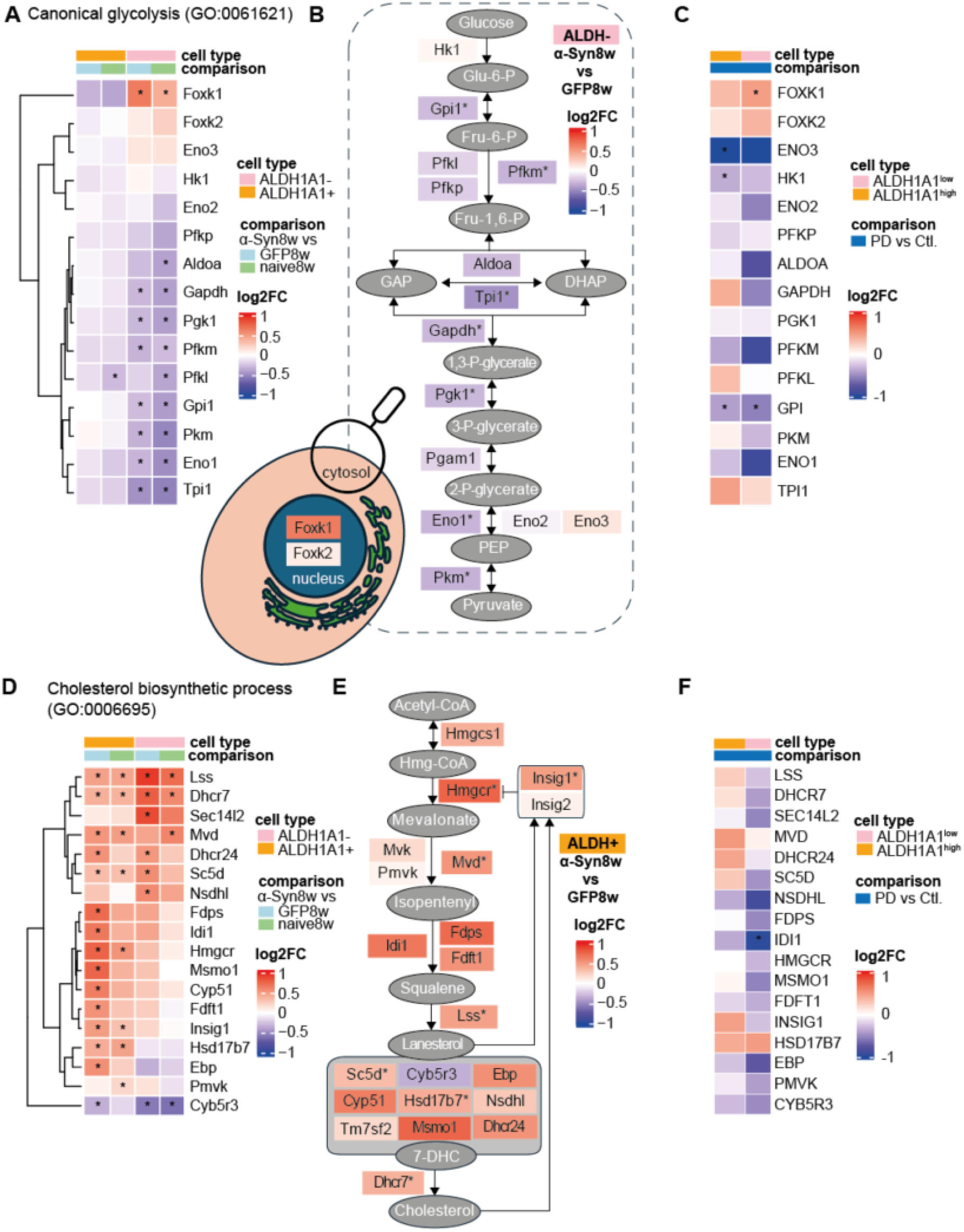
Subpopulation-specific disruption of glycolysis and cholesterol biosynthesis genes in response to αSyn overexpression. (A) Heatmap of log2FC of genes involved in ‘Canonical glycolysis’ (GO:0061621) in ALDH1A1^+^ and ALDH1A1^−^ SN DANs. * p<0.05 in the comparisons as indicated (αSyn8w vs GFP8w; αSyn8w vs naive). (B) Schematic of the glycolytic pathway, with enzymes coloured by log2FC in ALDH1A1^−^ SN DANs for the αSyn8w vs GFP8w comparison. Asterisks indicate genes with p<0.05 in ALDH1A1^−^ SN DANs in both αSyn8w vs control comparisons. (C) Heatmap of differential expression of glycolytic genes from A), calculated from snRNASeq data published by Kamath et al.^34^. Log2FC values are shown for PD vs. control comparisons in both ALDH1A1^high^ and ALDH1A1^low^ subpopulations. * p<0.05. (D) Heatmap of log2FC of genes involved in ‘Cholesterol biosynthetic process’ (GO:0006695) in ALDH1A1^+^ and ALDH1A1^−^ SN DANs. * p<0.05 in the comparisons as indicated (αSyn8w vs GFP8w; αSyn8w vs naive). (E) Schematic of cholesterol biosynthesis from acetyl-CoA to cholesterol, with enzymes coloured by log2FC in ALDH1A1^+^ SN DANs for the αSyn8w vs. GFP8w comparison. Asterisks indicate genes with p<0.05 in ALDH1A1^+^ SN DANs in both αSyn8w vs control comparisons. (F) Heatmap of differential expression of cholesterol biosynthesis genes from D), calculated from snRNASeq data published by ^34^. Log2FC values are shown for PD vs. control comparisons in both ALDH1A1^high^ and ALDH1A1^low^ subpopulations. * p<0.05.

In contrast to the suppression of bioenergetic pathways observed in ALDH1A1^−^ DANs, ALDH1A1^+^ DANs showed strong up-regulation of genes involved in cholesterol biosynthesis (Figure 6C), with these genes identified as LRGs. Indeed, several cholesterol biosynthesis genes were among the most highly dysregulated in ALDH1A1^−^ DANs (Figure 4E). This included up-regulation of the rate-limiting HMG-CoA reductase (*Hmgcr*). The cholesterol 24-hydroxylase *Cyp46a1*, a mediator of cholesterol efflux, was also down-regulated, indicating potential retention of cholesterol in the brain. Acetyl-CoA metabolism was also enriched in ALDH1A1^+^ DANs, along with more general sterol metabolism processes (Figure 5D). Together, this suggested a later metabolic shift toward sterol/cholesterol metabolism in ALDH1A1^+^ DANs.

To evaluate the translational relevance of these metabolic signatures, we re-analysed single-nuclei RNA-Seq data from human DANs in the SNpc of PD and control brains^34^. We re-classified Kamath *et al.*’s reported dopaminergic clusters into ALDH1A1^high^ and ALDH1A1^low^ subpopulations (Supp. Figure 2), and compared the expression of glycolysis- and cholesterol-related genes between PD and control samples in each sub-population. Glycolysis genes showed a general trend towards down-regulation in PD in both subpopulations (Figure 6C), resembling the suppression of glycolytic enzymes observed in ALDH1A1^−^ DANs in our mouse model. *FOXK1* was also up-regulated in both human subpopulations, suggesting a conserved pro-glycolytic transcriptional response. We also observed a trend towards selective up-regulation of cholesterol biosynthesis genes in human ALDH1A1^high^ DANs (Figure 6F), in alignment with the mouse data.

## DISCUSSION

Dopaminergic neurons (DANs) in the midbrain are molecularly, anatomically and functionally heterogenous, and this heterogeneity is increasingly recognized as an important factor shaping their responses to PD-associated stressors. Several studies have identified ALDH1A1^+^ DAN subpopulations in the midbrain^16,18,49^, but to our knowledge the spatially-resolved transcriptomic profiles of ALDH1A1+ and ALDH1A1^−^DANs, and their temporal responses to αSyn pathology, have not been thoroughly characterized *in situ*. Here, we mapped the spatiotemporal transcriptomic changes of ALDH1A1^+^ and ALDH1A1^−^ DAN subpopulations in the mouse midbrain following AAV-mediated αSyn overexpression.

Our data showed that these ALDH1A1^+^ and ALDH1A1^−^ DAN subpopulations are molecularly distinct. We identified robust transcriptomic differences between ALDH1A1^+^ and ALDH1A1^−^ naïve DANs that expanded on previously-reported ALDH1A1^+^ marker genes. Transcriptomic differences also revealed specialized cell-type-specific functions. Notably, ALDH1A1^−^ marker genes were enriched for various synaptic and neurotransmitter functions (including dopamine), and we also observed selective down-regulation of synaptic function genes in ALDH1A1^−^ DANs following αSyn overexpression. αSyn is normally enriched in presynaptic terminals^6,7^ and regulates synaptic function and neurotransmitter release through multiple mechanisms: it is involved in synaptic vesicle endocytosis^50^, regulates the organisation of synaptic vesicles^51^, and contributes to the assembly of the SNARE complex^52^. The synaptic ‘specialisation’ of ALDH1A1^−^ DANs may make them more sensitive to αSyn-induced synaptic stress in this model. Indeed, early synaptic dysfunction often precedes cell loss in PD^53^.

ALDH1A1^−^ DANs also displayed early bioenergetic dysfunction following αSyn over-expression, characterised by down-regulation of genes involved in glycolysis and other canonical ATP-producing pathways. Bioenergetic dysfunction is an established molecular mechanism implicated in PD. Within the glycolytic pathway, gene variants in *GAPDH*, the glycolytic rate-limiting enzyme, have been linked to an increased PD risk in men^54^. Similarly, a mutation in *PGK1*, the first ATP-producing enzyme in glycolysis and located within the *PARK12* susceptibility locus, is associated with parkinsonism^55^. We observed strong cell-type-specific down-regulation of genes spanning the entire glycolytic pathway, including *Gapdh* and *Pgk1*. Although we observed a similar trend in ALDH1A1^+^ DANs, changes were milder and not significant, suggesting a phenotype more resilient to αSyn-induced glycolytic suppression. Our data support strategies to boost glycolytic flux or stabilise mitochondrial ATP production. Terazosin, a PGK1 activator, has shown promising results in pre-clinical PD models^56–58^ and is now in Phase II clinical trials (NCT05109364: first posted 2021-11-05).

In contrast to the bioenergetic and synaptic down-regulation observed in ALDH1A1^−^SN DANS, we identified a selective up-regulation of genes enriched for acetyl-CoA and sterol, specifically cholesterol, metabolism, in ALDH1A1^+^ SN DANs. This included up-regulation of *Hmgcr*, the rate-limiting enzyme in cholesterol biosynthesis and an established target of statins^59^. Interestingly, up-regulation of cholesterol biosynthesis genes was accompanied by down-regulation of *Cyp46a1*, an enzyme that converts cholesterol into 24S-hydroxycholesterol, facilitating cholesterol efflux from the brain^60^. Together, enhanced cholesterol synthesis and reduced cholesterol efflux may suggest cholesterol retention in our model. The dysregulation of cholesterol metabolism raises the question around statin use and PD. However, studies have reported both a decreased and increased PD risk associated with statins^61^, and plasma cholesterol levels have been reported as both positive and negative risk factors for PD^62^. Whether cholesterol retention in the brain may be beneficial, compensatory, or further contribute to toxicity remains to be resolved. A potential dual role is illustrated by efavirenz, an anti-HIV medication and allosteric activator of *Cyp46a1*. While it has demonstrated neuroprotective properties in αSyn transgenic mice by reducing αSyn propagation^63^, it has also been shown to induce mitochondrial dysfunction and oxidative stress^64^, well-known PD mechanisms, and parkinsonism symptoms have recently been reported in people with chronic, virally-supressed HIV^65^. Our data suggest that any cholesterol-targeting medication in PD may have cell-type- and time/stage-specific effects, and these effects should be carefully considered.

Together, these findings support a model in which αSyn-associated pathology does not affect all midbrain DANs uniformly, but engages subpopulation-specific molecular pathways that may vary according to disease stage, anatomical location and model context. This distinction is important when interpreting ALDH1A1-defined subpopulations, as previous studies have reported model-dependent patterns of vulnerability, with ALDH1A1^+^ DANs preferentially affected in some contexts and ALDH1A1^−^ DANs in others. Within the timeframe analysed following αSyn overexpression, we observed significant reductions in TH^+^/ALDH1A1^−^ DANs in the SN at both 3- and 8-week timepoints, while TH^+^/ALDH1A1^+^ DANs showed only a subtle, non-significant decline at the later timepoint. This pattern aligns with a previous transgenic mouse study that showed greater vulnerability of ALDH1A1^−^ neurons to αSyn pathology^17^, and contrasts with findings from mitochondrial toxin models and human PD tissue, where ventral ALDH1A1^+^ DANs were under-represented in PD compared to controls^16,24,31^, implying differences in vulnerability of ALDH1A1-defined DAN subpopulations to key, PD-relevant stress conditions.

ALDH1A1 is proposed to confer neuroprotection by converting the highly reactive DOPAL metabolite to DOPAC, thereby catalyzing the metabolism of cytotoxic DA by-products^19–21^. In agreement, inhibition of ALDH1A1 leads to elevated DOPAL levels and has been demonstrated to promote αSyn aggregation^22,23,66^. To this end, previous studies have postulated that loss of ALDH1A1, either through decreased expression^17,24^, cleavage by asparagine endopeptidase (AEP)^67^, or inhibition by the fungicide benomyl^68^, may render these DANs susceptible to pathology. In addition, *Aldh1a1* knockout increases DA release and extracellular DA in the mouse striatum^69^, suggesting that DA release from ALDH1A1^+^ DANs may be restricted by ALDH1A1-mediated metabolism of DA. In PD therefore, as synaptic DA levels decline, ALDH1A1 expression may be reduced as a compensatory mechanism to increase DA release, potentially increasing their vulnerability. This would suggest that rescuing synaptic and bioenergetic function, down-regulated in ALDH1A1^−^ DANs at relatively early stages in the presented model, could be critical to prevent phenotypic change, or loss of either cell type. The relative preservation of ALDH1A1^+^ DANs in our study should not be interpreted as intrinsic resistance, however, as these neurons may be affected at later stages or under different pathological conditions. ALDH1A1 expression may be better viewed as one component of a broader molecular continuum, rather than as a standalone marker of vulnerability or resilience^46^. Indeed, heterogeneity even within ALDH1A1-defined subpopulations is already reported^34,70^, and other determinants of selective DAN vulnerability may include their elaborate axonal arborization, anatomical profile, dysregulated calcium loading, and high mitochondrial bioenergetic demands^71^.

This study highlights the value of spatial techniques to separate subtle yet distinct effects across DAN subpopulations *in situ*, with the ability to resolve meaningful functional differences potentially contributing to selective vulnerability. The addition of our findings to the literature suggests that distinct mechanisms may drive cell loss and adaptation in different models and at different stages of PD. As discussed above, ALDH1A1^−^ DANs in our model showed stronger early αSyn-induced synaptic and bioenergetic dysfunction, while the relative preservation of ALDH1A1⁺ DANs may reflect their lack of synaptic specialization and the protective properties of ALDH1A1 itself. In human PD, additional mechanisms likely co-exist to varying extents, or could emerge as pathology progresses. However, distinguishing these across cell types may be difficult in *post-mortem* tissue, where extensive cell loss has usually already occurred. Nevertheless, human data that we re-analysed here supported partial conservation of metabolic dysregulation.

## CONCLUSION

We here provide a comprehensive spatial and temporal characterization of mouse midbrain ALDH1A1^+^ and ALDH1A1^−^ DANs, revealing cell-type-specific transcriptomic responses to αSyn overexpression. Our findings suggest a model in which αSyn overexpression drives subpopulation-specific synaptic and metabolic stress, and adaptations that may underlie differential vulnerability. Future molecular therapeutic strategies should consider differential cell-type-specific responses across dopaminergic brain regions, model types, and disease stages.

## Acknowledgments

This work was funded by the Fulbright/RCSI StAR International PhD programme, an initiative of the Fulbright Commission and RCSI, University of Medicine & Health Sciences, and has received support from RCSI’s International Secondment Award programme, the Helmholtz Visiting Researcher Grant, and the FAWCO Foundation Education Awards programme. This publication has emanated from research supported in part by a research grant from Research Ireland under Grant Number 21/RC/10294_P2 and co-funded under the European Regional Development Fund and by FutureNeuro industry partners and Grant Numbers 21/FFP-A/9209 and 18/RI/5792. This study was supported by the EU Joint Programme – Neurodegenerative Disease Research and the Health Research Board [Grant number JPND-2023-2] under the 4DPD-Omics project. ÉBR was supported by a NeuroInsight Fellowship under the EU’s Horizon 2020 research and innovation programme under the Marie Skłodowska-Curie grant number 101034252. JVS was funded by Research Ireland through the Research Ireland Centre for Research Training in Genomics Data Science under grant number 18/CRT/6214. The authors acknowledge Heiko Dussmann, Ina Woods, Karen Conboy, Albert Sanfeliu, Alison Murphy and Sinead O’Sullivan for technical support.

## Author contributions

CJS performed experiments, analysed data and wrote the manuscript. ÉBR performed experiments. JVS, FN and KC analysed data. DCH, DAD and JHP provided supervision and secured funding. AU performed experiments, provided supervision and secured funding. NMCC analysed data, provided supervision, secured funding and wrote the manuscript. All authors read and approved the final manuscript.

## Abbreviations

AAV: adeno-associated virus;
ALDH1A1: aldehyde dehydrogenase 1A1;
AOI: area of interest;
αSyn: alpha-synuclein;
DAN(s): dopaminergic neuron(s);
DEG: differentially expressed gene;
DOPAL: 3,4-dihydroxyphenylacetaldehyde;
ERG: early responding gene;
LRG: late responding gene;
GFP: green fluorescent protein;
LMM: linear mixed model;
PD: Parkinson’s disease;
PEA: pathway enrichment analysis;
ROI: region of interest;
SN: substantia nigra;
SNCA: alpha-synuclein gene;
TH: tyrosine hydroxylase;
VTA: ventral tegmental area

## Legends for Supplementary Figures and Data

**Supplementary Figure 1: Comparison of differential expression in αSyn8w vs naïve and αSyn8w vs GFP8w comparisons.** Scatter plots showing log_2_FC of DEGs from αSyn8w vs naïve (x-axis) and αSyn8w vs GFP8w (y-axis) comparisons in (A) ALDH1A1^+^ and (B) ALDH1A1^−^ DANs. Genes with p<0.05 in one comparison are plotted in grey, and genes with p<0.05 in both conditions are plotted in purple, with their corresponding regression lines and correlation coefficients. Snca and EGFP (|log2FC|>5) were removed for visualization.

**Supplementary Figure 2: ALDH1A1 expression across dopamine neuron clusters from single-nucleus RNA-seq dataset generated by Kamath et al**^34^. Violin plots show normalized ALDH1A1 expression levels across the ten annotated DA neuron clusters, while the dot plot summarises the percentage of ALDH1A1-expressing nuclei and relative average expression within each population. We re-classified Kamath et al.’s clusters into ALDH1A1^high^ and ALDH1A1^low^ subpopulations, as indicated by the coloring of the cell Identity labels.

**Supplementary Data 1: Output of differential expression and pathway enrichment analyses in midbrain ALDH1A1^+^ vs. ALDH1A1^−^ DANs.**

**Supplementary Data 1a:** Differential expression output for naïve ALDH1A1+ vs naïve ALDH1A1– midbrain DANs (visualized in Figure 3A).

**Supplementary Data 1b:** Pathway enrichment output (GOBP) for midbrain ALDH1A1+ and ALDH1A1– marker genes (visualized in Figure 3C).

**Supplementary Data 2: Output of differential expression and pathway enrichment analysis in SN DANs.**

**Supplementary Data 2a:** Differential expression output for 1) ALDH1A1+ SN DANs – αSyn8w vs controls: αSyn8w vs naive and αSyn8w vs GFP8w, Figure 4E; 2) ALDH1A1– SN DANs – αSyn8w vs controls: αSyn8w vs naive and αSyn8w vs GFP8w, Figure 4F; and 3) ALDH1A1+ and ALDH1A1– SN DANs – αSyn3w vs αSyn8w.

**Supplementary Data 2b:** Pathway enrichment output (GOBP) for DEGs from 1) ALDH1A1+ SN DANs – αSyn8w vs controls; and 2) ALDH1A1– SN DANs – αSyn8w vs controls (Figure 4G)

**Supplementary Data 2c:** Pathway enrichment output (GOBP) for ERGs and LRGs from both ALDH1A1+ and ALDH1A1– SN DANs (Figure 5D)

## REFERENCES

1 Steinmetz, J. D. et al. Global, regional, and national burden of disorders affecting the nervous system, 1990&#x2013;2021: a systematic analysis for the Global Burden of Disease Study 2021. The Lancet Neurology 23, 344–381, doi:10.1016/S1474-4422(24)00038-3 (2024).

2 Dehay, B. et al. Targeting α-synuclein for treatment of Parkinson’s disease: mechanistic and therapeutic considerations. Lancet Neurol 14, 855–866, doi:10.1016/s1474-4422(15)00006-x (2015).

3 Farrow, S. L., Cooper, A. A. & O’Sullivan, J. M. Redefining the hypotheses driving Parkinson’s diseases research. NPJ Parkinsons Dis 8, 45, doi:10.1038/s41531-022-00307-w (2022).

4 Spillantini, M. G. et al. Alpha-synuclein in Lewy bodies. Nature 388, 839–840, doi:10.1038/42166 (1997).

5 Baba, M. et al. Aggregation of alpha-synuclein in Lewy bodies of sporadic Parkinson’s disease and dementia with Lewy bodies. Am J Pathol 152, 879–884 (1998).

6 Maroteaux, L., Campanelli, J. T. & Scheller, R. H. Synuclein: a neuron-specific protein localized to the nucleus and presynaptic nerve terminal. J Neurosci 8, 2804–2815, doi:10.1523/JNEUROSCI.08-08-02804.1988 (1988).

7 Kahle, P. J. et al. Subcellular Localization of Wild-Type and Parkinson’s Disease-Associated Mutant α-Synuclein in Human and Transgenic Mouse Brain. The Journal of Neuroscience 20, 6365–6373, doi:10.1523/jneurosci.20-17-06365.2000 (2000).

8 Sharma, M. & Burre, J. alpha-Synuclein in synaptic function and dysfunction. Trends Neurosci 46, 153–166, doi:10.1016/j.tins.2022.11.007 (2023).

9 Simuni, T. et al. A biological definition of neuronal &#x3b1;-synuclein disease: towards an integrated staging system for research. The Lancet Neurology 23, 178–190, doi:10.1016/S1474-4422(23)00405-2 (2024).

10 Polymeropoulos, M. H. et al. Mutation in the &#x3b1;-Synuclein Gene Identified in Families with Parkinson’s Disease. Science 276, 2045–2047, doi:doi:10.1126/science.276.5321.2045 (1997).

11 Singleton, A. B. et al. alpha-Synuclein locus triplication causes Parkinson’s disease. Science 302, 841, doi:10.1126/science.1090278 (2003).

12 Garritsen, O., van Battum, E. Y., Grossouw, L. M. & Pasterkamp, R. J. Development, wiring and function of dopamine neuron subtypes. Nat Rev Neurosci 24, 134–152, doi:10.1038/s41583-022-00669-3 (2023).

13 Damier, P., Hirsch, E. C., Agid, Y. & Graybiel, A. M. The substantia nigra of the human brain. II. Patterns of loss of dopamine-containing neurons in Parkinson’s disease. Brain 122 (Pt 8), 1437–1448, doi:10.1093/brain/122.8.1437 (1999).

14 Fearnley, J. M. & Lees, A. J. Ageing and Parkinson’s disease: substantia nigra regional selectivity. Brain 114 (Pt 5), 2283–2301, doi:10.1093/brain/114.5.2283 (1991).

15 Maingay, M., Romero-Ramos, M., Carta, M. & Kirik, D. Ventral tegmental area dopamine neurons are resistant to human mutant alpha-synuclein overexpression. Neurobiol Dis 23, 522–532, doi:10.1016/j.nbd.2006.04.007 (2006).

16 Poulin, J. F. et al. Defining midbrain dopaminergic neuron diversity by single-cell gene expression profiling. Cell Rep 9, 930–943, doi:10.1016/j.celrep.2014.10.008 (2014).

17 Liu, G. et al. Aldehyde dehydrogenase 1 defines and protects a nigrostriatal dopaminergic neuron subpopulation. J Clin Invest 124, 3032–3046, doi:10.1172/jci72176 (2014).

18 Pereira Luppi, M., et al. Sox6 expression distinguishes dorsally and ventrally biased dopamine neurons in the substantia nigra with distinctive properties and embryonic origins. Cell Reports 37, 109975, doi:10.1016/j.celrep.2021.109975 (2021).

19 Marchitti, S. A., Deitrich, R. A. & Vasiliou, V. Neurotoxicity and Metabolism of the Catecholamine-Derived 3,4-Dihydroxyphenylacetaldehyde and 3,4-Dihydroxyphenylglycolaldehyde: The Role of Aldehyde Dehydrogenase. Pharmacological Reviews 59, 125–150, doi:10.1124/pr.59.2.1 (2007).

20 Rees, J. N., Florang, V. R., Anderson, D. G. & Doorn, J. A. Lipid peroxidation products inhibit dopamine catabolism yielding aberrant levels of a reactive intermediate. Chem Res Toxicol 20, 1536–1542, doi:10.1021/tx700248y (2007).

21 Goldstein, D. S. et al. Determinants of buildup of the toxic dopamine metabolite DOPAL in Parkinson’s disease. J Neurochem 126, 591–603, doi:10.1111/jnc.12345 (2013).

22 Burke, W. J. et al. Aggregation of alpha-synuclein by DOPAL, the monoamine oxidase metabolite of dopamine. Acta Neuropathol 115, 193–203, doi:10.1007/s00401-007-0303-9 (2008).

23 Masato, A. et al. DOPAL initiates αSynuclein-dependent impaired proteostasis and degeneration of neuronal projections in Parkinson’s disease. NPJ Parkinsons Dis 9, 42, doi:10.1038/s41531-023-00485-1 (2023).

24 Galter, D., Buervenich, S., Carmine, A., Anvret, M. & Olson, L. ALDH1 mRNA: presence in human dopamine neurons and decreases in substantia nigra in Parkinson’s disease and in the ventral tegmental area in schizophrenia. Neurobiology of Disease 14, 637–647, doi:10.1016/j.nbd.2003.09.001 (2003).

25 Vila, M. et al. α-Synuclein Up-Regulation in Substantia Nigra Dopaminergic Neurons Following Administration of the Parkinsonian Toxin MPTP. Journal of Neurochemistry 74, 721–729, 10.1046/j.1471-4159.2000.740721.x (2000).

26 Kowall, N. W. et al. MPTP induces alpha-synuclein aggregation in the substantia nigra of baboons. NeuroReport 11, 211–213 (2000).

27 Van Den Berge, N. & Ulusoy, A. Animal models of brain-first and body-first Parkinson’s disease. Neurobiol Dis 163, 105599, doi:10.1016/j.nbd.2021.105599 (2022).

28 Cai, H., Liu, G., Sun, L. & Ding, J. Aldehyde Dehydrogenase 1 making molecular inroads into the differential vulnerability of nigrostriatal dopaminergic neuron subtypes in Parkinson’s disease. Translational Neurodegeneration 3, 27, doi:10.1186/2047-9158-3-27 (2014).

29 Giguère, N., Burke Nanni, S. & Trudeau, L. E. On Cell Loss and Selective Vulnerability of Neuronal Populations in Parkinson’s Disease. Front Neurol 9, 455, doi:10.3389/fneur.2018.00455 (2018).

30 Tang, L. et al. A primate nigrostriatal atlas of neuronal vulnerability and resilience in a model of Parkinson’s disease. Nat Commun 14, 7497, doi:10.1038/s41467-023-43213-2 (2023).

31 Rey, N. et al. Calbindin and Girk2/Aldh1a1 define resilient vs vulnerable dopaminergic neurons in a primate Parkinson’s disease model. npj Parkinsons Dis. 10, doi:10.1038/s41531-024-00777-0 (2024).

32 Goralski, T. M. et al. Spatial transcriptomics reveals molecular dysfunction associated with cortical Lewy pathology. Nat Commun 15, 2642, doi:10.1038/s41467-024-47027-8 (2024).

33 Ya, D. et al. Application of spatial transcriptome technologies to neurological diseases. Front Cell Dev Biol 11, 1142923, doi:10.3389/fcell.2023.1142923 (2023).

34 Kamath, T. et al. Single-cell genomic profiling of human dopamine neurons identifies a population that selectively degenerates in Parkinson’s disease. Nature Neuroscience 25, 588–595, doi:10.1038/s41593-022-01061-1 (2022).

35 Zhang, T. et al. Brain-wide alterations revealed by spatial transcriptomics and proteomics in COVID-19 infection. Nat Aging 4, 1598–1618, doi:10.1038/s43587-024-00730-z (2024).

36 Ulusoy, A. et al. Brain-to-stomach transfer of alpha-synuclein via vagal preganglionic projections. Acta Neuropathol 133, 381–393, doi:10.1007/s00401-016-1661-y (2017).

37 Migdalska-Richards, A. et al. The L444P Gba1 mutation enhances alpha-synuclein induced loss of nigral dopaminergic neurons in mice. Brain 140, 2706–2721, doi:10.1093/brain/awx221 (2017).

38 Milanese, C. et al. Activation of the DNA damage response in vivo in synucleinopathy models of Parkinson’s disease. Cell Death Dis 9, 818, doi:10.1038/s41419-018-0848-7 (2018).

39 Dinnon, K. H., 3rd et al. SARS-CoV-2 infection produces chronic pulmonary epithelial and immune cell dysfunction with fibrosis in mice. Sci Transl Med 14, eabo5070, doi:10.1126/scitranslmed.abo5070 (2022).

40 Yu, G., Wang, L. G., Han, Y. & He, Q. Y. clusterProfiler: an R package for comparing biological themes among gene clusters. OMICS 16, 284–287, doi:10.1089/omi.2011.0118 (2012).

41 Love, M. I., Huber, W. & Anders, S. Moderated estimation of fold change and dispersion for RNA-seq data with DESeq2. Genome Biol 15, 550, doi:10.1186/s13059-014-0550-8 (2014).

42 Ulusoy, A., Decressac, M., Kirik, D. & Bjorklund, A. Viral vector-mediated overexpression of alpha-synuclein as a progressive model of Parkinson’s disease. Prog Brain Res 184, 89–111, doi:10.1016/S0079-6123(10)84005-1 (2010).

43 McCaffery, P. & Drager, U. C. High levels of a retinoic acid-generating dehydrogenase in the meso-telencephalic dopamine system. Proc Natl Acad Sci U S A 91, 7772–7776, doi:10.1073/pnas.91.16.7772 (1994).

44 La Manno, G. et al. Molecular Diversity of Midbrain Development in Mouse, Human, and Stem Cells. Cell 167, 566–580.e519, doi:10.1016/j.cell.2016.09.027 (2016).

45 Carmichael, K. et al. Function and Regulation of ALDH1A1-Positive Nigrostriatal Dopaminergic Neurons in Motor Control and Parkinson’s Disease. Frontiers in Neural Circuits 15 (2021).

46 Yaghmaeian Salmani, B., et al. Transcriptomic atlas of midbrain dopamine neurons uncovers differential vulnerability in a Parkinsonism lesion model. Elife 12, doi:10.7554/eLife.89482 (2024).

47 Chang, C. H., Lim, K. L. & Foo, J. N. Synaptic Vesicle Glycoprotein 2C: a role in Parkinson’s disease. Front Cell Neurosci 18, 1437144, doi:10.3389/fncel.2024.1437144 (2024).

48 Sukonina, V. et al. FOXK1 and FOXK2 regulate aerobic glycolysis. Nature 566, 279–283, doi:10.1038/s41586-019-0900-5 (2019).

49 Tiklová, K. et al. Single-cell RNA sequencing reveals midbrain dopamine neuron diversity emerging during mouse brain development. Nat Commun 10, 581, doi:10.1038/s41467-019-08453-1 (2019).

50 Vargas, K. J. et al. Synucleins regulate the kinetics of synaptic vesicle endocytosis. J Neurosci 34, 9364–9376, doi:10.1523/JNEUROSCI.4787-13.2014 (2014).

51 Zaltieri, M. et al. alpha-synuclein and synapsin III cooperatively regulate synaptic function in dopamine neurons. J Cell Sci 128, 2231–2243, doi:10.1242/jcs.157867 (2015).

52 Burre, J., Sharma, M. & Sudhof, T. C. alpha-Synuclein assembles into higher-order multimers upon membrane binding to promote SNARE complex formation. Proc Natl Acad Sci U S A 111, E4274–4283, doi:10.1073/pnas.1416598111 (2014).

53 Bridi, J. C. & Hirth, F. Mechanisms of alpha-Synuclein Induced Synaptopathy in Parkinson’s Disease. Front Neurosci 12, 80, doi:10.3389/fnins.2018.00080 (2018).

54 Ping, Z. et al. GAPDH rs1136666 SNP indicates a high risk of Parkinson’s disease. Neurosci Lett 685, 55–62, doi:10.1016/j.neulet.2018.06.011 (2018).

55 Guimaraes, T. G. et al. X-Linked Levodopa-Responsive Parkinsonism-Epilepsy Syndrome: A Novel PGK1 Mutation and Literature Review. Mov Disord Clin Pract 11, 556–566, doi:10.1002/mdc3.13992 (2024).

56 Cai, R. et al. Enhancing glycolysis attenuates Parkinson’s disease progression in models and clinical databases. Journal of Clinical Investigation 129, 4539–4549, doi:10.1172/JCI129987 (2019).

57 Weber, M. A. et al. Glycolysis-enhancing alpha(1)-adrenergic antagonists modify cognitive symptoms related to Parkinson’s disease. NPJ Parkinsons Dis 9, 32, doi:10.1038/s41531-023-00477-1 (2023).

58 Kokotos, A. C. et al. Phosphoglycerate kinase is a central leverage point in Parkinson’s disease-driven neuronal metabolic deficits. Sci Adv 10, eadn6016, doi:10.1126/sciadv.adn6016 (2024).

59 Endo, A. The discovery and development of HMG-CoA reductase inhibitors. J Lipid Res 33, 1569–1582 (1992).

60 Pikuleva, I. A. & Cartier, N. Cholesterol Hydroxylating Cytochrome P450 46A1: From Mechanisms of Action to Clinical Applications. Front Aging Neurosci 13, 696778, doi:10.3389/fnagi.2021.696778 (2021).

61 Al-Kuraishy, H. M. et al. Pros and cons for statins use and risk of Parkinson’s disease: An updated perspective. Pharmacol Res Perspect 11, e01063, doi:10.1002/prp2.1063 (2023).

62 Alza, N. P., Iglesias González, P. A., Conde, M. A., Uranga, R. M. & Salvador, G. A. Lipids at the Crossroad of α-Synuclein Function and Dysfunction: Biological and Pathological Implications. Front Cell Neurosci 13 (2019).

63 Kim, J. B. et al. Artificial intelligence-driven drug repositioning uncovers efavirenz as a modulator of alpha-synuclein propagation: Implications in Parkinson’s disease. Biomed Pharmacother 174, 116442, doi:10.1016/j.biopha.2024.116442 (2024).

64 Lanman, T., Letendre, S., Ma, Q., Bang, A. & Ellis, R. CNS Neurotoxicity of Antiretrovirals. J Neuroimmune Pharmacol 16, 130–143, doi:10.1007/s11481-019-09886-7 (2021).

65 Shorer, E. F. et al. Parkinsonism in people with virally suppressed HIV. Lancet HIV 13, e194–e206, doi:10.1016/S2352-3018(25)00262-0 (2026).

66 Masařík, M. et al. Sensitive Electrochemical Detection of Native and Aggregated α-Synuclein Protein Involved in Parkinson’s Disease. Electroanalysis 16, 1172–1181, 10.1002/elan.200403009 (2004).

67 Nie, S. et al. Sox6 and ALDH1A1 Truncation by Asparagine Endopeptidase Defines Selective Neuronal Vulnerability in Parkinson’s Disease. Adv Sci (Weinh) 12, e2409477, doi:10.1002/advs.202409477 (2025).

68 Fitzmaurice, A. G. et al. Aldehyde dehydrogenase inhibition as a pathogenic mechanism in Parkinson disease. Proc Natl Acad Sci U S A 110, 636–641, doi:10.1073/pnas.1220399110 (2013).

69 Sgobio, C. et al. Aldehyde dehydrogenase 1-positive nigrostriatal dopaminergic fibers exhibit distinct projection pattern and dopamine release dynamics at mouse dorsal striatum. Sci Rep 7, 5283, doi:10.1038/s41598-017-05598-1 (2017).

70 Azcorra, M. et al. Unique functional responses differentially map onto genetic subtypes of dopamine neurons. Nature Neuroscience 26, 1762–1774, doi:10.1038/s41593-023-01401-9 (2023).

71 Surmeier, D. J., Obeso, J. A. & Halliday, G. M. Selective neuronal vulnerability in Parkinson disease. Nat Rev Neurosci 18, 101–113, doi:10.1038/nrn.2016.178 (2017).

